# Expression and characterization of the complete cyanophage genome PP in the heterologous host *Synechococcus elongatus* PCC 7942

**DOI:** 10.1101/2024.07.23.604706

**Authors:** Guorui Li, Jia Feng, Xiaofei Zhu, Yujie Chai, Tao Sun, Jianlan Jiang

**Affiliations:** School of Chemical Engineering & Technology, Tianjin University, Tianjin 300072, P.R. China; Key Laboratory of Systems Bioengineering (Ministry of Education), Tianjin University, Tianjin 300072, P. R. China; Frontier Science Center for Synthetic Biology (Ministry of Education), Tianjin University, Tianjin 300072, P.R. China; Center for Biosafety Research and Strategy, Tianjin University, Tianjin 300072, P.R. China

**Keywords:** cyanophage, genome design and assembly, cyanobacteria, inducible switches, transcriptomics

## Abstract

Cyanophages are considered a promising biological management option for treating cyanobacterial blooms. Broadening the host range of cyanophages and/or shortening the lysis cycle by designing and synthesizing artificial cyanophages are potential strategies to enhance their effectiveness and efficiency. However, the rescue of artificial cyanophage genomes remains unexplored. In this study, we achieved the integration of a full-length cyanophage genome, PP, which originally infects *Plectonema boryanum* FACHB-240, into the model cyanobacterium *Synechococcus elongatus* PCC 7942. Since the integration of these large fragments (∼42 kb) into cyanobacteria depended on conjugation via *Escherichia coli*, the toxic open reading frames (ORFs) of PP to *E. coli* were first identified, leading to the identification of toxic ORF6, ORF11, and ORF22. The original PP genome was then rearranged, and the three toxic ORFs were controlled using a tandem induction switch. The full length of the PP genome was integrated into the genome of *S. elongatus* PCC 7942 via two rounds of homologous recombination.

Interestingly, compared to the control strain, the integration of the PP genome decreased photosynthesis and carbon fixation in *S. elongatus* PCC 7942, exhibiting cyanophage-like behavior. Transcriptomic analysis revealed that 32 of the 41 ORFs of the PP genome were transcribed in *S. elongatus* PCC 7942, significantly altering the energy metabolism and carbon fixation pathways. These influences were further demonstrated using metabolomics. This study provides a comprehensive approach for the artificial design and integration of cyanophage genomes in cyanobacteria, laying the foundation for their real rescue in the future.

## 1. Introduction

Photosynthetic cyanobacteria play an important ecological role in terms of global carbon sinks and oxygen sources. However, harmful cyanobacterial blooms block sunlight from other organisms and consume large amounts of oxygen and nutrients, causing damage to the hydrosphere. Blooming cyanobacteria also produce and release a wide range of cyanotoxins, which can pose an indirect or direct threat to the health of humans and animals including liver damage, some immunotoxicity, reproductive toxicity, or neurotoxicity by contaminating drinking water (1). The most famous case of the cyanobacterial toxin Cylindrospermopsin, known as the Palmetto Mystery Disease, occurred in Australia in 1979, whose residual toxin in drinking water caused by a local algaecide led to the poisoning of over a hundred children with gastroenteritis (2). In the U.S., more than 300 medical visits were reported related to harmful cyanobacteria between 2017-2019, involving skin, gastrointestinal, respiratory and neurological conditions (3). In this case, efficient strategies to control the abundance and burst frequency of blooming cyanobacteria are essential for environmental safety and human health.

Approaches for blooming cyanobacteria removal usually involve prevention and control. In addition, physical methods (*i.e.*, mechanical salvage, physical aeration), ultrasonic treatment, chemical methods (*i.e.*, algal removers or flocculants), and biological methods (*i.e.*, biological algaecides and bioactive materials) have been explored or utilized in different studies (4). Nevertheless, these strategies face challenges such as being time-consuming, demanding, costly, or causing secondary pollution. Thus, exploration of fast, efficient and economical treatment solutions is necessary. Cyanophage is a kind of planktonic virus in the environment with cyanobacteria as the main host, which can rapidly infect and lyse cyanobacteria in suitable environment (5). Given its ability to specifically lyse cyanobacterial cells, cyanophage has been considered as an important biological regulator of cyanobacterial population density and a promising biological management option for treating cyanobacterial blooms (6). Nevertheless, their infection is highly host-specific, which depends on the binding of receptor-binding proteins to host surface receptors (7). This leads to the fact that each cyanophage has its strict host range, making it difficult to control with only one or a few natural cyanophages.

To address the issue, broadening the host range of cyanophages and/or shortening the lysis cycle by designing and synthesizing artificial cyanophages via synthetic biology are promising strategies. Previously, our group realized the design and assembly of two cyanophage genomes, i.e., PP (8) and A-4L (9) in *Saccharomyces cerevisiae*, but facing challenges in their rescue or heterogenous expression in cyanobacteria. Transferring large DNA fragments into cyanobacteria generally depends on the conjugation method mediated by *Escherichia coli*, but full assembly of PP and A-4L was found lethal to *E. coli*. Thus, we believe the challenges are mainly caused by the toxic genes containing by cyanophages. To explore the rescue method of artificially constructed cyanophage genome, we focused on the cyanophage PP originally infecting the cyanobacterium *Plectonema boryanum* FACHB-240 in this study. Briefly, the potential toxic genes or clusters to *E. coli* on genome of PP were systematically evaluated and identified first. As genetic manipulation method for *P. boryanum* FACHB-240 has not been fully established, we aimed to transfer the artificial cyanophage genome into model cyanobacterium *Synechococcus elongatus* PCC 7942 (hereafter PCC 7942), which is suitable for transformation of large DNA fragments up to 107.9 kb (10) and offers a similar genetic background to *P. boryanum* FACHB-240. Via genome rearrangement and driving the toxic gene clusters of PP using strictly inducible promoters, we successfully integrated the full PP cyanophage genome (including the beginning and end terminal repeat sequences) into the chromosome of PCC 7942 with two rounds’ homologous recombination. Interestingly, we found the significant growth defect and physiological changes in engineered PCC 7942 with PP genome after induction, suggesting the expression of the heterogenous cyanophage. To explore the interactions between the PP genome and PCC 7942 host, we performed transcriptional analysis and elucidated the related genes or pathways. The work here provides new insights for assembly and rescue cyanophages in the future.

## 2. Material and methods

### 2.1. Strains, competent cells, plasmids, and culture conditions

All strains, competent cells and plasmids constructed or used in this study are listed in **Table S1**. *E. coli* was cultured in LB medium with applicable antibiotics (i.e., 25 mg/mL chloramphenicol, 50 mg/mL kanamycin) at 37°C according to the different plasmids carried, respectively. *E. coli* used for toxicity gene validation experiments were cultured at 37°C in LB medium with different antibiotics (25 mg/mL chloramphenicol, 50 mg/mL kanamycin) and containing inducible promoter inducers (theophylline at a final concentration of 2 mM, and IPTG at a final concentration of 1 mM), according to the different plasmids carried by them, respectively. *S. cerevisiae* BY4741 was cultured in YPD (Yeast Extract Peptone Dextrose) medium at 30°C, 200 rpm. Yeast strains were cultured in synthetic complete medium without histidine/uracil (SC–H/U), depending on the different plasmids carried.

All the PCC 7942 strains were cultured at 37°C with a light density of μmol photons/(m^2^·s). 7942-sm was cultured in BG11 medium containing 25 μg/mL streptomycin. The constructed strain 7942-halfPP was cultured in BG11 liquid medium containing 25 μg/mL streptomycin and 25 μg/mL chloramphenicol. 7942-PP was cultured in BG11 liquid medium containing 25 μg/mL streptomycin, 25 μg/mL chloramphenicol and 25 μg/mL kanamycin. The control strain 7942-sm and the experimental strain 7942-PP used for transcriptome analysis were additionally supplemented with inducible promoter inducer (theophylline and IPTG with a final concentration of 2mM and 1 mM, respectively).

### 2.2. Yeast assembly

*S. cerevisiae* BY4741 competent cells were prepared as described (11). Briefly, after overnight culture, yeast cells were added to 5 mL of fresh medium and harvested when OD_600_ reached 0.5. Yeast competent cells were prepared with 0.1 M LiOAc and stored briefly in an ice box. DNA fragments with about 60-100 bp overlaps and linearized vector were co-transformed into 100 μL competent cells referring to the LiAc/SS carrier DNA/PEG method. The cells were then spread onto corresponding solid nutrient-deficient media and cultured at 30°C for 2 days to identify positive clones. Yeast colony PCR analysis was performed according to a modified method (12). Recombinant plasmids extracted from yeast using TIAN prep Mini Plasmid Kit (TIANGEN BIOTECH, Beijing, China) and High Pure BAC DNA Mini Kit (Magen Biotechnology, Guangdong, China) were transferred into *E. coli* competent cells.

### 2.3. Transformation of *Synechococcus elongatus* PCC 7942

The plasmids PSynPP4-8 (37, 449bp) and PSynPP1-3I3 (30, 185bp) that were transformed to PCC 7942 in this study were both large fragments, thus the best way to transform them would be by triparental conjugative transfer rather than natural transformation. The PSynPP4-8 plasmid and PSynPP1-3I3 plasmid were transferred from *E. coli* to PCC 7942 via conjunction. *E. coli* containing the target plasmid and *E. coli* containing helper plasmids including pRL 443 and pRL 623 were cultured to logarithmic phase. 2 mL of the bacterial solution was taken and washed three times with LB medium, and then resuspended in 100 μL of LB medium, respectively. Since the medium of the cyanobacteria in this study always contained different types of antibiotics, a total of 5 mL of PCC 7942 cells at logarithmic phase needed to be centrifuged and washed three times with BG11 medium and resuspended in 0.2 mL of BG 11 medium. All the above suspensions were mixed and incubated at 37°C and 100 μmol photons/(m^2^·s) for 60 min. The mixture was inoculated on BG11 plus 5% LB (vol/vol) agar plates, with a nitrocellulose membrane with pore diameter of 0.45 mm, at 30 °C for 24 hours under μmol photons/(m^2^·s). The membrane was then transferred to a medium containing corresponding antibiotics (50 μg/mL streptomycin and 50 μg/mL chloramphenicol for 7942-halfPP; 50 μg/mL streptomycin, 50 μg/mL chloramphenicol and 50 μg/mL kanamycin for 7942-PP) and cultured on a BG11 agar plate for 5 to 7 days to screen stable transformants. Cyanobacterial transformants were verified by colony PCR and subjected to Sanger sequencing.

### 2.4. Phenotype analysis

Fresh seed solutions (7942-sm, 7942-PP) were inoculated into 20 mL of fresh liquid medium containing the required antibiotics and inducers mentioned above. The optical density (OD) at 750 nm and the full absorption spectrum of the liquid cultures were measured by using a Varioskan LUX multifunctional microplate reader (Thermo Fisher Scientific, Waltham, MA, USA). The initial OD_750_ of the liquid culture was 0.05 and was measured every 12 h. The full absorption spectrum and OJIP of the strain cells were measured at 96 h, using 1 mL samples by the AquaPen AP 110/C Handheld PAM Fluorometer (FluorCam, Drásov, Czech Republic). Each strain was repeated by three biological parallel samples.

### 2.5. Transcriptomic analysis

Transcriptomic analyses of 7942-sm and 7942-PP cultured for 96 h with inducers were performed to assess the heterologous expression of the full-length genome of PP cyanophage in PCC 7942 and its effect on heterologous hosts. Transcriptomic analysis was performed using the RNA-sequencing method (RNA-seq) in AZENTA (Suzhou, China). Three parallel replicates were included for each sample. After the quality assessment of raw reads obtained by transcriptome sequencing, low-quality reads were pre-processed to obtain clean reads. The HTSeq software (v.0.6.1) was used to evaluate gene expression levels, based on the FPKM metric. Deseq2 (v.1.6.3) within the Bioconductor software program was then used to identify DEGs, based on comparisons of experimental and control groups. DEGs were identified with fold changes of >1.5 and a P value of <0.05.

### 2.6. Quantitative Real-Time PCR (qRT-PCR)

Cyanobacterial cultures in liquid medium were incubated for 96 hours (with a cell culture volume*OD750 = 5), followed by centrifugation, immediate RNA extraction, and freezing with liquid nitrogen. The RNA from each sample was quantified and used to synthesize cDNA using the Hiscript II Q RT SuperMix for qPCR (Vazyme, Nanjing, China). The obtained cDNA was diluted 100-fold and used as the template for qRT-PCR. The qRT-PCR mixtures were prepared with the ChamQ Universal SYBR qPCR Master Mix (Vazyme, Nanjing, China) according to the manufacturer’s instructions. The qRT-PCRs were performed using the StepOne real-time PCR system (Applied Biosystems, CA, USA). The *rnpB* gene, encoding the RNase P subunit B, was used as a housekeeping gene. Each sample was analyzed in three technical replicates. The qRT-PCR data were analyzed using the StepOne software program (Applied Biosystems) and the 2^-ΔΔC^ method (13). Genes were selected to validate RNA-seq expression data with qRT-PCR. The correlations between RNA-seq and qRT-PCR were assessed using Pearson correlation coefficients with Excel. These correlation coefficients were used to identify reliable RNA-seq expression data.

## 3. Results

### 3.1. At least 3 genes toxic to *E. coli* existed in the PP genome

The natural cyanophage PP genome is a 42,480 bp (GenBank: KF598865) linear double-stranded DNA with 222 bp terminal repeats at both ends, encoding a total of 41 open reading frames (ORFs), 17 of which are functionally annotated (14, 15) **(Table S2)**. As mentioned above, the complete assembly of the PP genome in *E. coli* is key point as it will mediate the transfer of PP genome to PCC 7942 via conjugation. Based on the natural cyanophage PP genome, we divided it into 8 DNA fragments (i.e., PP1, PP2…, PP8) for chemical synthesis and artificial assembly based on a shuttle vectors pRS426 (containing 2μ and ColE1 origin of replication) between *S. cerevisiae* and *E. coli*. Each of them was about 5 kb in length and carried homology arms of about 300 bp in neighboring fragments. Interestingly, though all the 8 fragments were successfully assembled in *S. cerevisiae* (i.e., pRSPP1, pRSPP2…, pRSPP8; **Table S1**), we found that plasmids pRSPP1, pRSPP2, and pRSPP3 respectively containing the first three DNA fragments (*i.e.*, PP1, PP2 and PP3) could not be transformed into *E. coli* (**Figure 1A**). Meanwhile, consistent with our previous report (8), the whole assembled plasmid pRSPP4-8 containing 5 parts from PP4 to PP8 could survive in *E. coli*. However, its combination with any of the PP1, PP2 or PP3 fragment led to the failed clone in *E. coli*, suggesting toxic genes existed in PP1, PP2, and PP3 (according to the location of 223∼16,487 bp of the cyanophage PP genome). To identify the potential toxic ORFs in PP1, PP2 and PP3, they were sub-divided into 9 fragments (1∼2 kb) named with PP1-F1, PP1-F2, PP1-F3, PP1-F4, PP2-F5, PP2-F6, PP2-F7, PP3-F8, and PP3-F9 (the according plasmids were pRSPP1-F1, pRSPP1-F2…, pRSPP3-F8; **Table S1**). Surprisingly, all of them could survive in *E. coli* (**Figure 1B**). In this case, we deduced that toxic genes may just existed in the neighboring regions of the 9 sub-fragments, which were disrupted due to the artificial separation.

**Figure 1.**
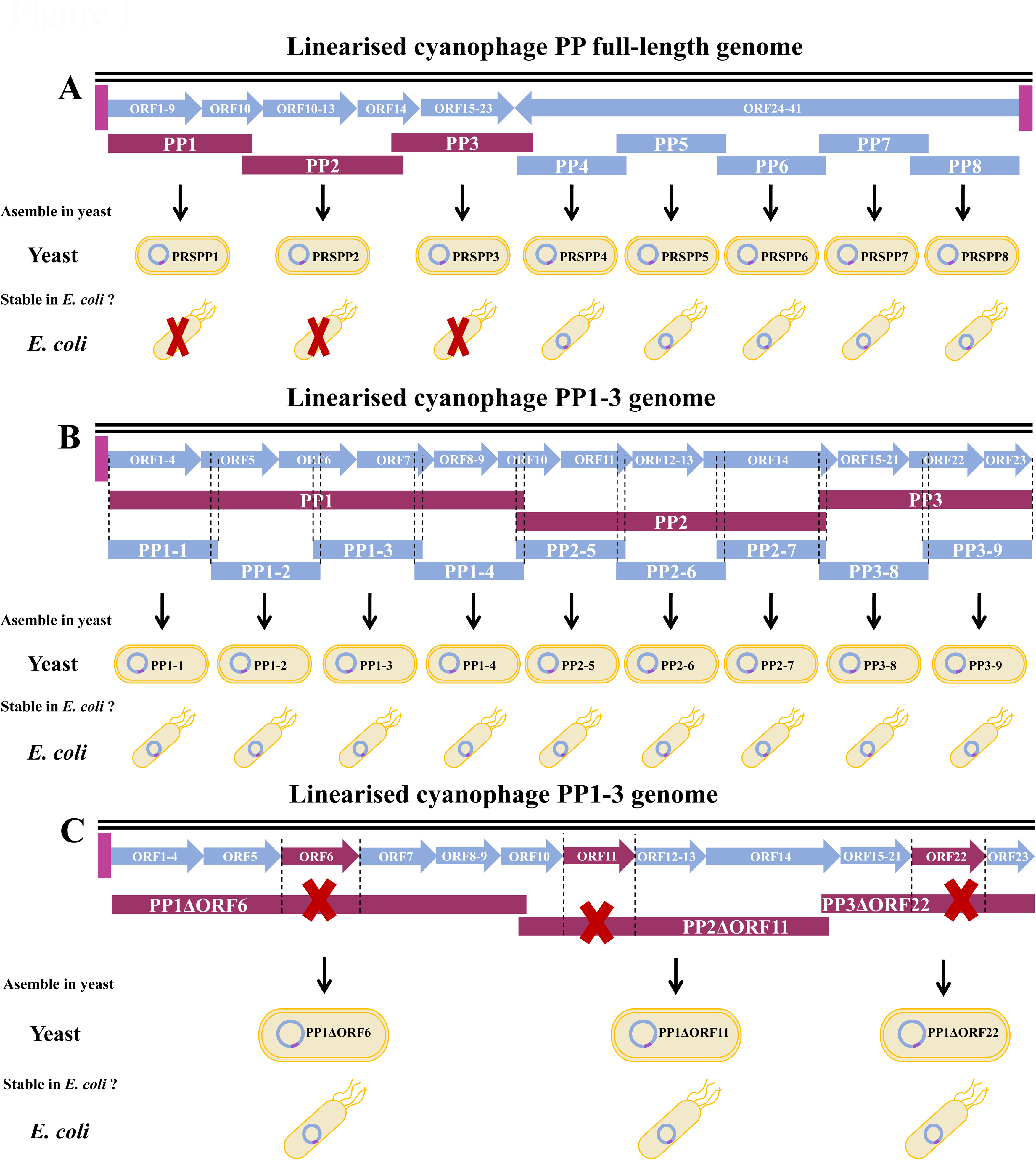
Schematic and the concluded results of the toxic gene screen of PP genome. (**A**) Identification of the large toxic fragments of PP genome. (**B**) Identification of the toxic fragments of PP1, PP2 and PP3. (**C**) Identification of the large toxic ORFs of PP1, PP2 and PP3.

We then focused on the 7 ORFs including ORF5, ORF6, ORF7, ORF10, ORF11, ORF14, and ORF22, which existed in the neighboring regions of the 9 sub-fragments. Further, we respectively deleted these ORFs from the original large fragments PP1, PP2, PP3 or PP1-3, assembling 14 plasmids both using the high-copy pRS426 and single-copy pLS0 (16) for *E. coli* (the according plasmids were pRSPP1ΔORF5, pLSPP1ΔORF5…, pRSPP3ΔORF22 and pLSPP3ΔORF22; **Table S1**). As a result (**Figure 1C)**, constructers could survive using plasmids with deletion of ORF6 from PP1, ORF11 from PP2 and ORF22 from PP3, which was irrelevant to plasmid copy number. Not surprisingly, assembled plasmid pRSPP1-3Δ3 and pLSPP1-3Δ3 (containing PP1, PP2 and PP3) lacking ORF6, ORF11 and ORF22 could also survive in *E. coli* (**Figure 1C)**, suggesting these three ORFs were toxic to *E. coli*.

### 3.2. Toxic gene control via strictly inducible promoters enabled the stable existence of PP1-3 in *E. coli*

As mentioned above, pRSPP1-3Δ3 and pLSPP1-3Δ3 lacking ORF6, ORF11 and ORF22 maintained stable in *E. coli*. To create a complete assembly of PP1, PP2 and PP3 with all ORFs, we aimed to rearrange the original location of each ORF and control the toxic ORFs via strictly inducible switches. To ensure the universality of the inducible switches between *E. coli* and PCC 7942 as well as the minimum leakage, IPTG-inducible promoter and theophylline-inducible riboswitch were selected and used in tandem (17, 18). Firstly, pLSPP1-3I3 was assembled in *S. cerevisiae*, in which toxic ORF6, ORF11 and ORF22 (original promoters were removed and designed as polycistron) were controlled by AND switch responding to IPTG and theophylline **(Figure 2A**; **Table S1)**. The positive plasmid extracted from yeast transformants was then transformed into competent *E. coli.* Excitingly, pLSPP1-3I3 could be transferred into *E. coli* normally, suggesting the feasibility of our strategy. To further verify the functionality of toxic genes, we compared the growth of *E. coli* containing pLSPP1-3I3 with or without induction. As expected, addition of the two inducers (1 mM IPTG and 2 mM theophylline) led to the cell death of *E. coli* **(Figure 2B, C)**. However, assembled plasmid pRSPP1-8Δ3 and pLSPP1-8I3 (containing 8 parts from PP1 to PP8) lacking ORF6, ORF11 and ORF22 or controlling them via inducible swtiches still led to cell death of *E. coli*, indicating interactions existed between ORFs located in PP1-3 and in PP4-8 **(Figure 2D**; **Table S1)**. Alternatively, we tried to divided the whole PP genome into two parts, *i.e.*, PP1-3 and PP 4-8, transferring them into the PCC 7942 mediated by *E. coli* via two rounds.

**Figure 2.**
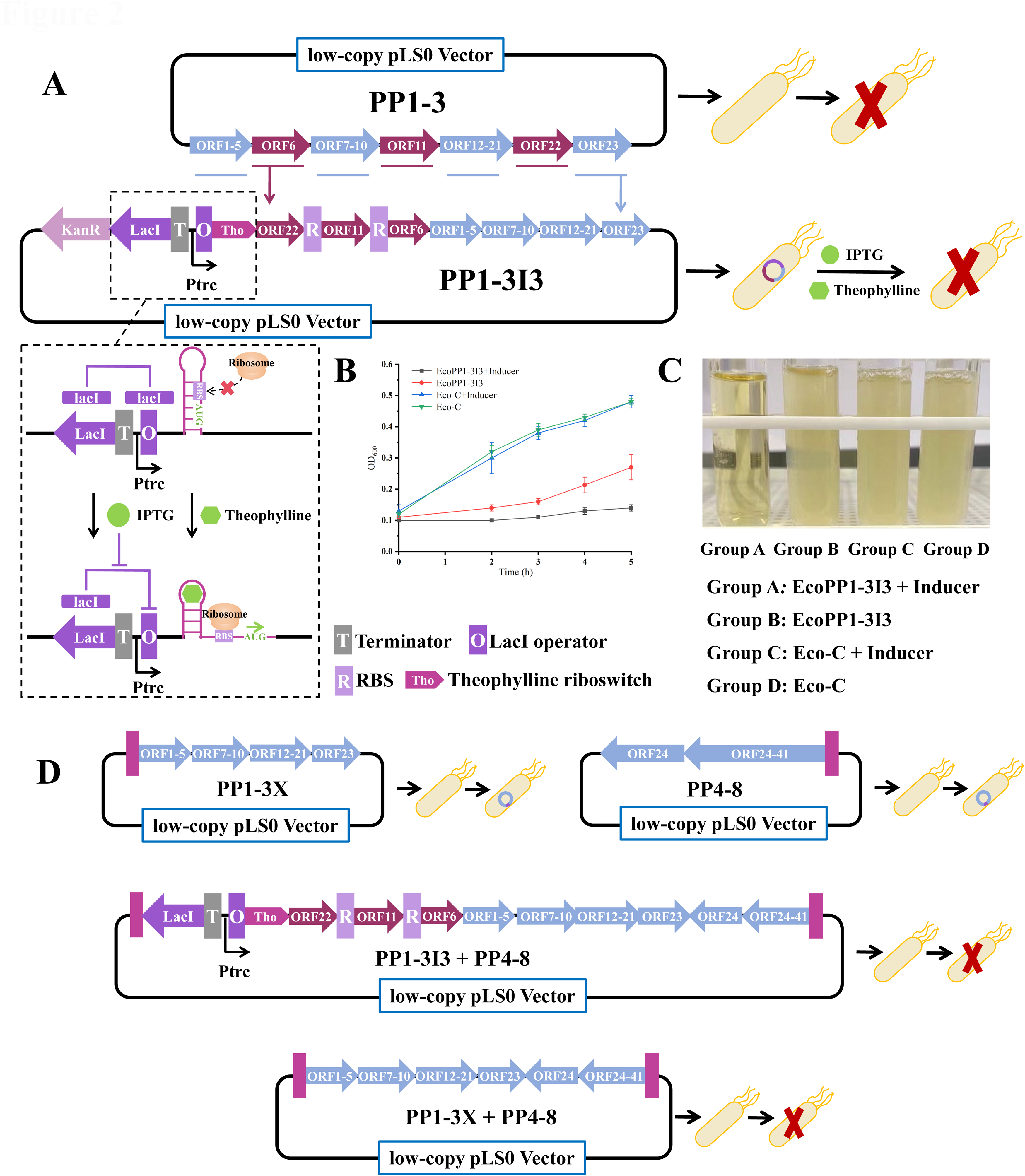
The strategy and validation of controlling toxic genes using the tandem induction switch. (**A**) The green rectangle is an artificially selected homology arm of about one thousand bp for subsequent homology substitution experiments. Red arrows are ORFs that can express lethal toxicity to *E. coli* with a single gene. Each of the three toxic ORFs was ligated with the low-copy vector pLS0, transformed into *E. coli*, and cultured with the corresponding resistant LB solid medium, and no transformants appeared. (**B**) Growth curves of Eco-C and EcoPP1-3I3 with or without inducers. (**C**) Pictures of Eco-C and EcoPP1-3I3 with or without inducers. (**D**) The assembly and transfer of a full PP genome in *E. coli*.

### 3.3. Rational design of the two parts of PP genome realized its successful integration into genome of PCC 7942 via two rounds’ homologous recombination

PCC 7942 is a well-studied model cyanobacterium, which has been demonstrated feasible for accepting large DNA fragments of >100 kb via conjugation (10). As we failed to constructed a stable plasmid containing a complete PP genome in *E. coli*, we aimed to integrated the PP genome into the genome of PCC 7942 via two rounds’ double-crossover homologous recombination. To ensure the successful integration and exclude false single-crossover homologous recombination (19), we adapted following designs based on the low-copy pLSPP4-8 to construct pSynPP4-8 (**Figure 3A**): i) two homologous arms targeting NSI site for recombination in PCC 7942 were added flanking the assembled fragments of PP genome; ii) additional recombinase λRed from pKD46 (20) was utilized to enhance the homologous recombination; iii) counter-selection marker *rpsL* (21) was added outside the homologous recombination region to exclude single-crossover recombination (a streptomycin-tolerant strain 7942-sm was created in this study and false integration of *rpsL* would made 7942-sm streptomycin-sensitive again).

**Figure 3.**
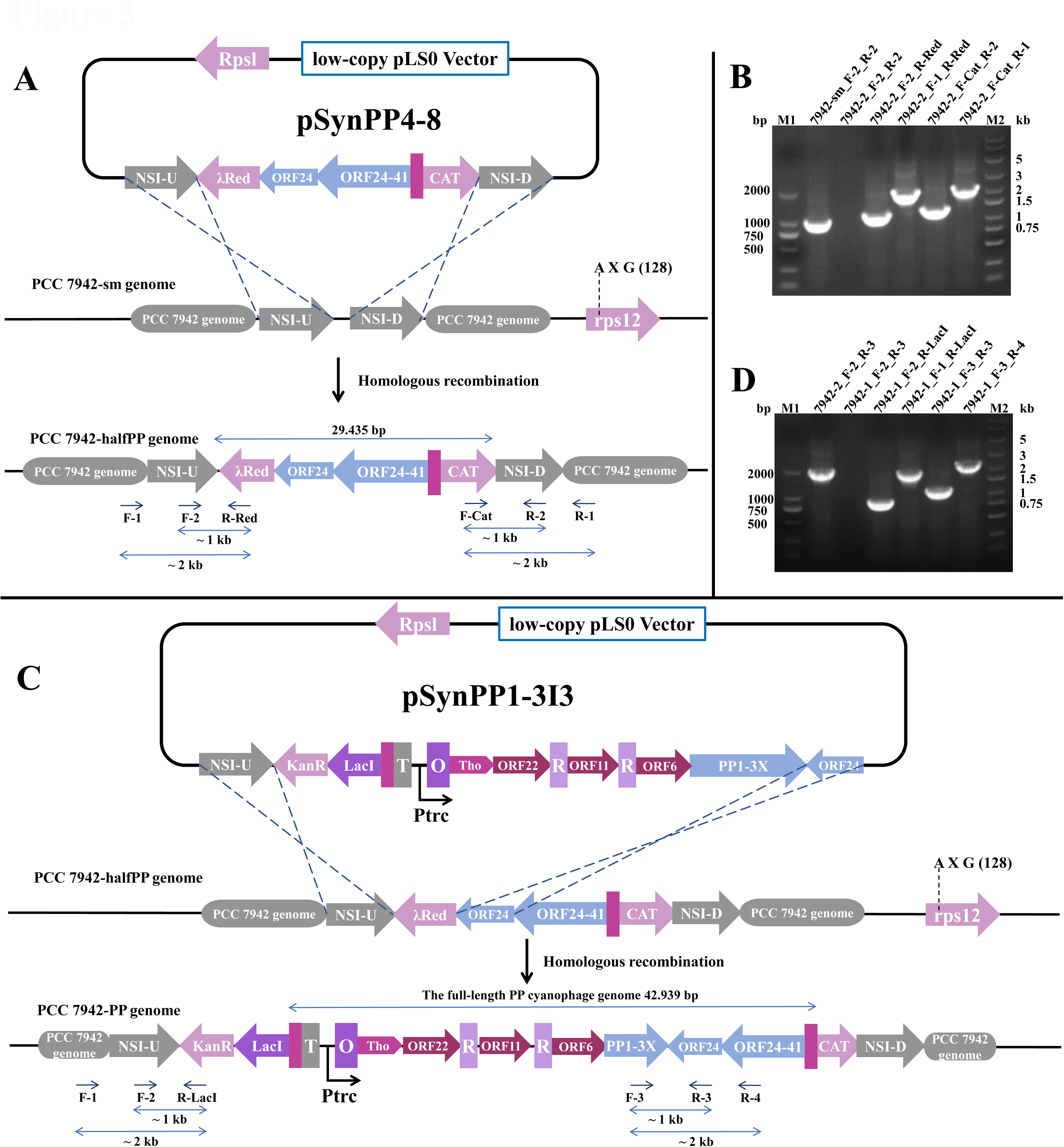
The strategy and validation of integrating the full-length genome of PP cyanophage into the genome of PCC 7942. RpsL is a gene encoding the ribosomal protein S12, which is a target protein for streptomycin action on the ribosome. When the RpsL gene is present in 7942-sm, streptomycin resistance takes on a cryptic shape and streptomycin sensitivity is a dominant trait. Therefore, we used the RpsL gene as a reverse screening marker to prevent single exchange during gene integration. λRed: The cyanophage lambda-derived Red recombination system. It is used to improve the accuracy and success of homologous recombination. CAT: Chloramphenicol acetyltransferase, inactivates chloramphenicol by acetylating it. ORF24: Starting part of PP4. (**A**) PSynPP4-8 plasmid for integration to the 7942-sm genome at NSI. Identification strategy of the 7942-halfPP genome after the first homologous recombination with the required primers. (**B**) The results of homologous recombination were identified by PCR. The template strains and primer information of the PCR products are labelled above the gel image. M1: DL 2000, M2: 250 bp. (**C**) The PSynPP1-3I3 plasmid used for integration into the 7942-halfPP genome at PP4-8. Identification strategy of the 7942-PP genome after the second homologous recombination with the required primers. (**D**) The results of homologous recombination were identified by PCR. The template strains and primer information of the PCR products are labelled above the gel image. M1: DL 2000, M2: 250 bp.

The correctly identified *E. coli* transformants were transformed into 7942-sm (a streptomycin-tolerant strain of PCC 7942), leading to the strain 7942-halfPP. As shown in **Figure 3B**, the genomic integration of the 5 fragments PP4-8 was confirmed by PCR. Similarly, pSynPP1-3I3 was constructed for the second round of transformation based on pLSPP1-3I3. Detailly, pSynPP1-3I3 contained PP1-3 with toxic ORF6, ORF11 and ORF22 cassettes controlled by inducible switches, positive selection maker (kanamycin-resistant cassette, Km^R^), two homologous arms respectively targeting upstream of NSI site and partial of the PP4 (**Figure 3C**). Finally, the correct integration of pSynPP1-3I3 resulted in the insertion of PP1-3 between upstream NSI and PP4 (**Figure 3C**), leading to the strain 7942-PP. The second round of integration was confirmed by PCR (**Figure 3D**). Through the above strategy, we achieved the construction of a complete PP cyanophage genome including its repeat sequences of the first and the last ends in the non-host cyanobacterium PCC 7942.

### 3.4. Heterologous expression of the full-length genome of PP cyanophage exhibited multiple influences on physiology of PCC 7942

To evaluate the effect of PP genome cyanophage on PCC 7942, we simultaneously compared the growth curves, photosynthetic and carbon fixation capability of the control strain 7942-sm and engineered strain 7942-PP. Under the same light intensity and temperature conditions, we found that the growth rate of 7942-PP was slower than that of 7942-sm **(Figure 4A, 4B)**, suggesting that the expression of the PP cyanophage genome influences the growth of PCC 7942. In addition, though the addition of inducers or not had almost no effect on the growth of 7942-sm, the logarithmic phase of 7942-PP was delayed by 2-3 days after induction compared to that of non-inducer-added 7942-PP. The above results indicated that both the toxic genes controlled by the inducible promoter and other non-toxic genes affected the growth of PCC 7942 to some extent. On the other hand, we measured their absorption spectrum of 7942-sm and 7942-PP with the same biomass (both with OD_750_ = 0.55) with inducers **(Figure 4C)**. The absorption peaks of chlorophyll a and chlorophyll b were found to be significantly reduced in 7942-PP. This suggests that heterologous expression of the PP cyanophage genome affected the photosynthetic system of 7942-PP. Consistently, the chlorophyll fluorescence induction kinetics curve (OJIP) of 7942-sm and 7942-PP suggested that the PP cyanophage genome severely affected the PQ and electron transfer on the PSII receptor side of the photosynthetic system ofc considering the increased S_m_ and N value. **(Figure 4D)**. Finally, we found that content of sucrose and glycogen in 7942-PP was significantly lower than that in 7942-sm **(Figure 4E**, **4F)**. As these metabolites were the main products from carbon fixation, we deduced expression of the PP genome also affected the carbon fixation of 7942-PP. All these results suggested that genes of PP genome successfully expressed in 7942-PP and exerted similar influences like a real cyanophage to some extent.

**Figure 4.**
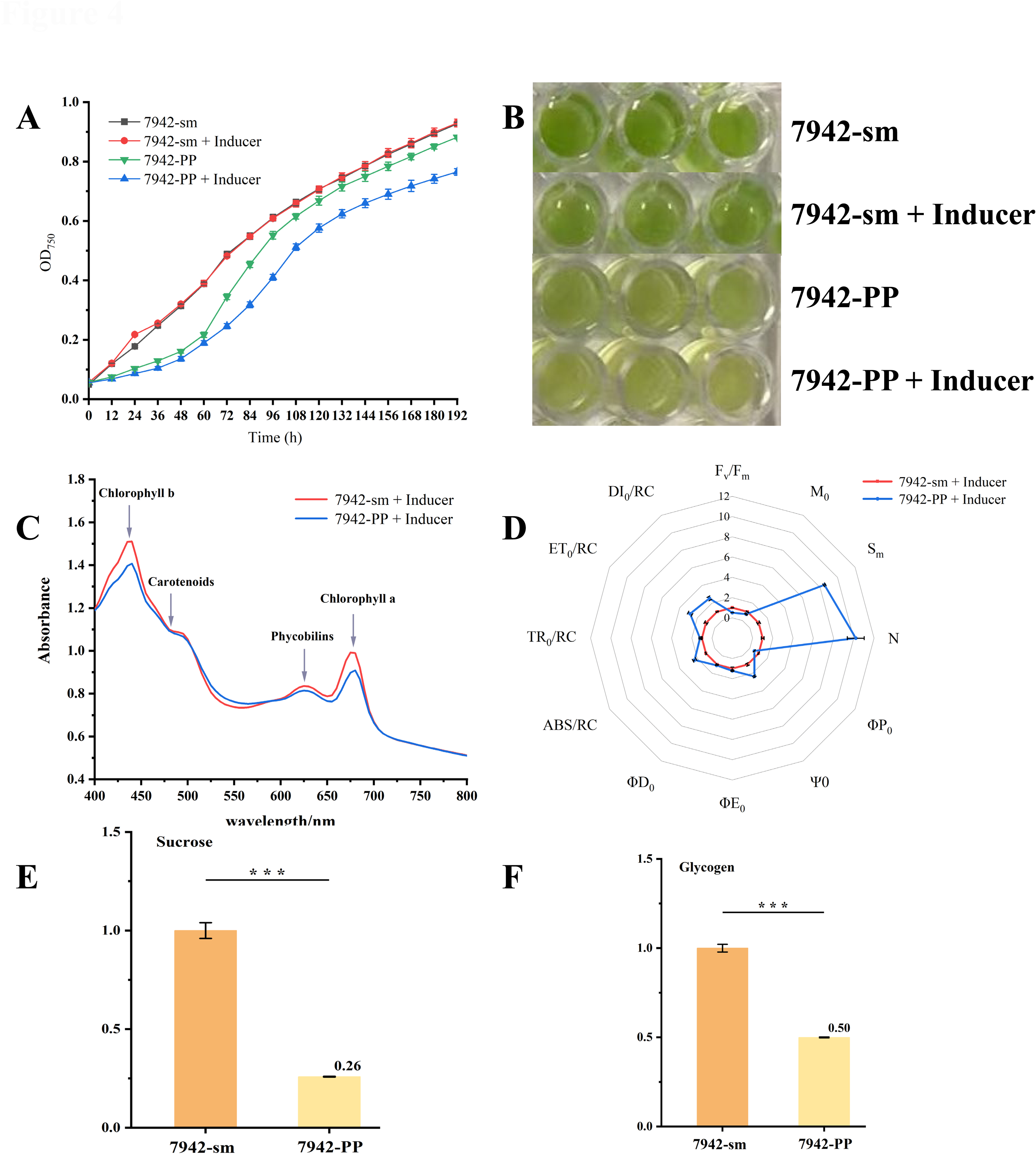
Physiological comparations of 7942-sm and 7942-PP. (**A**) Growth patterns of 7942-sm and 7942-PP with or without inducers. (**B**) Pictures of 7942-sm and 7942-PP with or without inducers. (**C**) Absorption spectrum of 7942-sm and 7942-PP after induction. (**D**) OJIP curve of 7942-sm and 7942-PP after induction. (**E**) Sucrose content of 7942-sm and 7942-PP after induction. (**F**) Glycogen content of 7942-sm and 7942-PP after induction.

### 3.5. Comparative transcriptome and metabolome analysis revealing the global effect of PP genome on PCC 7942

To further evaluate the global effects of PP genome on 7942-PP, transcriptional analysis of 7942-sm and 7942-PP cultured for 96 h with inducers was performed.

#### 3.5.1. Expression and function of cyanophage PP genes

FPKM (Fragments Per Kilo bases per Million reads) was measured for 41 ORFs that were predicted in the cyanophage PP genome. Excitingly, 34 of them showed detectable expressions in the transcriptional level, suggesting that PCC 7942 provided similar genetic background to the original host *P. boryanum* FACHB-240 (**Table S3**). For the remaining seven ORFs, including the tightly controlled ORF6, several factors could account for their behavior: i) these predicted ORFs may not be real; ii) their promoters may not be recognized by the heterologous host PCC 7942; or iii) their expression levels may be too low to reach the detection limit. Among the detected ORFs, ten are associated with nucleotide synthesis and DNA replication, six with amino acid synthesis and ribosome translation, and four with carbohydrate metabolism.

Detailly, ORF1 and ORF2 encode the DNA polymerase sliding clamp subunit (PCNA homolog). It plays an important coordinating role in DNA replication and DNA damage repair (22). This gene was first reported to be present in the phage genome. ORFs 5-7 encode PurL and ORFs 8-10 encode PurF. These genes are used to increase nucleotide synthesis during infection and facilitate the viral genome replication (23, 24). In addition, ORFs12-14, ORF17, and ORF39 all encode genes related to protein synthesis. ORF24 encodes 2-keto-3-deoxy-6-phosphogluconate aldolase, an enzyme specific to the ED (Entner-Doudoroff) pathway (25). This suggested that cyanophage PP promotes the ED pathway in the host to produce more NADPH (26). ORF35 and ORF36 encode the Gnd, which has been shown to be used by phages to regulate PPP (Pentose phosphate pathway) in cyanobacteria (23). The 6-phosphogluconate dehydrogenase encoded by the *gnd* effectively promotes PPP in the host, and we found that these genes were also present in the genome of PCC 7942, and their expression was significantly elevated. The expressions of these ORFs may explain the physiological changes we observed in the last section.

#### 3.5.2. Effect of heterologous expression of cyanophage PP gene on PCC 7942

Further, we analyzed the differentially expressed genes in 7942-PP compared to that in 7942-sm. Generally, using a 2-fold change as a cutoff, 667 of 2665 expressed genes were identified as differentially expressed genes (DEGs), including 290 down-regulated and 377 up-regulated genes (**Figure 5**, **S1A**). According to pathway enrichment, the enriched DEGs were mainly involved in Carbon metabolism, Carbon fixation in photosynthetic organisms, Biosynthesis of amino acids and Nucleotide metabolism (**Figure S1B**).

**Figure 5.**
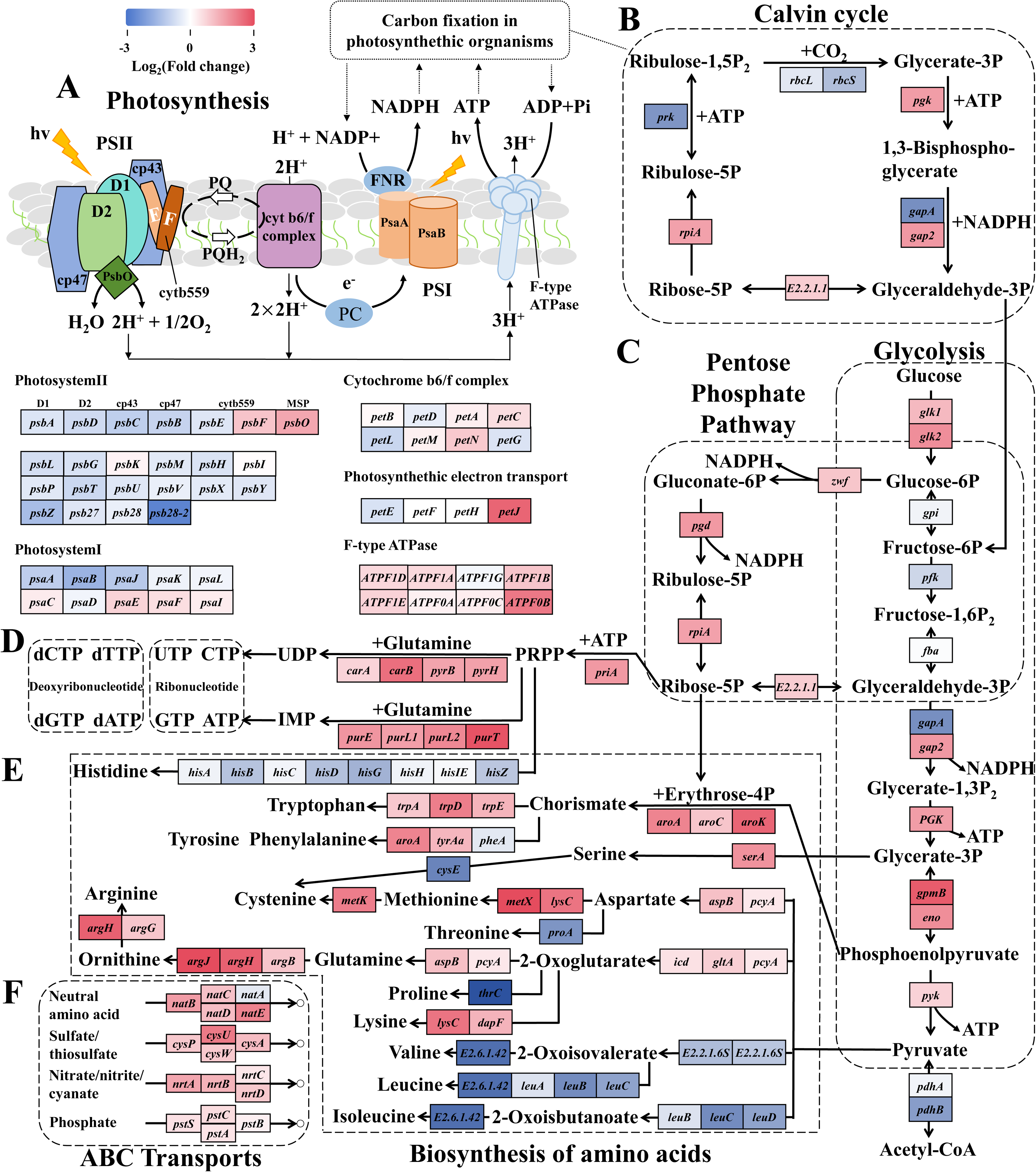
Response of PCC 7942 to heterologous expression of cyanophage PP genome. (**A**) Volcano map of differentially expressed genes. Red dots indicate up-regulation and blue dots indicate down-regulation. The abscissa represents the logarithm of the fold change of gene expression, and the ordinate represents the statistical significance of the change in gene expression. Differentially expressed genes (DEGs) were screened based on a fold change > 2 and p-value < 0.05. (**B**) KEGG pathway-enrichment scatter plot under normal culture conditions. The abscissa represents the Rich factor. The larger the Rich factor, the greater the enrichment degree. The ordinate represents the pathway name. The size of each dot indicates the number of DEGs in this pathway, and the color of each dot corresponds to different p-values. The smaller the p-value, the more significant the enrichment.

In the PCC 7942 genome, the expression of 19 genes associated with photosystem II was down-regulated, and the expression of 6 genes associated with photosystem I was down-regulated, with almost all of the genes used to synthesize photosystem proteins down-regulated (**Figure 5A**). In the PCC 7942 genome, the expression of 19 genes associated with photosystem II and 6 genes associated with photosystem I were down-regulated, with almost all of the genes used to synthesize photosystem proteins down-regulated, such as P680 reaction center D1/D2 protein, CP43/CP47 chlorophyll apoprotein, in photosystem II and P700 chlorophyll an apoprotein A1/A2 in photosystem I. These proteins play a critical role in the photosynthetic system, which suggests that the products expressed by the cyanophage PP genome severely affect the photosynthetic process in cyanobacteria. It is hypothesized that the cyanophage reduces the consumption of other amino acids for the synthesis of the relevant proteins. It is noteworthy that *psbO*, which is associated with the assembly of light reaction centers, showed a significant up-regulation, probably due to the weakening of photosynthesis, which led to a resistance mechanism generated by the cyanobacteria themselves. *petF* and *petJ*, as related genes associated with electron transport, were both significantly upregulated. *petF* encodes ferredoxin, an important component of the photosynthetic electron transport chain involved in the transfer of electrons to oxidoreductases and reduction of NADP^+^ to NADPH (27), which well explained our results in section 3.4. Although both chlorophyll a and chlorophyll b were reduced in 7942-PP **(Figure 4C)**, and both *F_V_/F_M_* and φ*_Po_*, representing maximum quantum yield for primary photochemistry, were decreased, absorption flux per RC and parameters related to electron transfer efficiency were substantially increased **(Figure 4D)**. This suggested that although the cyanophage PP genome leads to reduced synthesis of chlorophyll and photosynthetic proteins in cyanobacteria, energy supply is ensured by increasing the efficiency of electron transfer in the photosynthetic system. This is in contrast to the results of studies on marine cyanophages, which encode homologs of the core peptides D1 and D2 of the photosystem II reaction center in order to maintain the photosynthetic activity of the host and thus obtain sufficient energy for its own replication (28, 29). Different from the cyanophages with longer latency to infect, the latency of virulent cyanophage PP is extremely short, so it may not obtain energy by influencing the host to synthesize more photosynthesis proteins, but rather, it directly reduced the synthesis of photosynthesis proteins of the host in order to complete its own replication (30). The expression of 17 genes related to NADPH and ATP production was up-regulated in the photosynthetic system. Normally, these ATP and NADPH generated by photosynthesis go to participate in the dark reaction to convert CO_2_ to organic carbon (Calvin–Benson–Bassham cycle; CBB cycle). However, the expression of *rbcS* and *rbcL* genes encoding ribulose-bisphosphate carboxylase (Rubisco) was significantly down-regulated in the CBB cycle **(Figure 5B)**. The significant downregulation of *rbcS* and *rbcL* expression directly inhibits the CBB cycle, which saved more energy for replication of cyanophages (31). Similarly, reactions involving glyceraldehyde 3-phosphate dehydrogenase (phosphorylating), encoded by *gapA*, require the consumption of NADPH, and reactions involving phosphoribulokinase, encoded by *prk*, require the consumption of ATP. Their down-regulation allowed phagosomes to recruit these remaining NADPH and ATP to meet their nucleotide and metabolic requirements (32).

Other genes upregulated in the CBB cycle also play important roles in the PPP **(Figure 5C)**. The rate-limiting enzyme of the PPP is glucose-6-phosphate 1-dehydrogenase, and the significant up-regulation of *zwf*, which encodes it, together with other important genes in the cycle, suggests that the pentose phosphate cycle is promoted. Meanwhile, 6-phosphogluconate dehydrogenase encoded by ORF35 and ORF36 in the cyanophage PP genome further promoted PPP. CBB cycle and PPP are two pathways with opposite effects. The CBB cycle consumes NADPH and ATP to reduce CO_2_ to produce hexose, and the PPP utilizes glucose to produce CO_2_ and NADPH. Therefore, we speculated that the products expressed by the cyanophage PP genome direct the carbon flux from the CBB cycle to the PPP by inhibiting CO_2_ fixation in cyanobacteria, thereby obtaining a large amount of NADPH and ATP to meet the metabolic demands and energy supply for additional cyanophage genome replication. On the other hand, we could find up-regulation of *glk1*, *glk2*, *zwf* and down-regulation of *gpi*, *pfk* and *fba* in glycolysis, suggesting that more glucose was phosphorylated to Glucose 6-phosphate and directed to participate more in the PPP rather than the Embden-Meyerhof pathway (EMP) **(Figure 5C)**. At the same time glycolysis of core module involving three-carbon compounds was promoted, suggesting that glyceraldehyde 3-phosphate generated either by PPP or EMP was accelerated in the reaction, leading to the generation of more ATP with NADPH and large amounts of pyruvate. Interestingly, we found the *eda* gene encoding 2-keto-3-deoxy-6-phosphogluconate aldolase in the cyanophage PP genome. This is an enzyme specific to the ED pathway. The ED-introduced strain was found to consume glucose 1.5-fold faster than the parental strain (33). The ED pathway is characterized by the rapid acquisition of pyruvate from glucose in only 4 steps, as compared to 10 steps in the EMP pathway. Compared to EMP, the ED pathway produces less ATP and more NADPH for the same amount of glucose consumed, a trend consistent with the previous results (34). We proposed that phage-augmented NADPH production fuels deoxynucleotide biosynthesis for phage replication. Further, the expression of the genes encoding the PDH enzyme family were all down-regulated, indicating that the oxidative decarboxylation of pyruvate was inhibited. Pyruvate, as the end product of the glycolytic pathway, is oxidized to produce acetyl-CoA, but also participates in many important physiological reactions, such as for amino acid synthesis **(Figure 5E)**. Pyruvate can be used to synthesize a wide range of amino acids, but we found that the synthesis of proline, threonine, and branched-chain amino acids was severely inhibited, while the synthesis of other amino acids, such as glutamine, arginine and lysine, was significantly promoted. Glycerate 3-phosphate and phosphoenolpyruvate in the glycolysis of core module involving three-carbon compounds can be used to synthesize tryptophan, tyrosine, phenylalanine and cysteine. The synthesis of all these amino acids was promoted.

Ribose 5-phosphate, which participates in the CBB cycle and the PPP, is also important for the synthesis of synthetic amino acids. Ribose 5-phosphate is converted to phosphoribosyl pyrophosphate (PRPP) by ribose-phosphate pyrophosphokinase encoded by the *priA* gene, a reaction that is significantly facilitated **(Figure 5D)**. PRPP can be used to synthesize tryptophan. However, the genes that consume PRPP to synthesize tryptophan are all significantly down-regulated, presumably to preserve sufficient raw material for nucleotide synthesis. This is because PRPP is an extremely important precursor for the synthesis of purine and pyrimidine nucleotides. As can be seen from **Figure 5D**, the genes associated with the synthesis of deoxyribonucleotide and nucleotide are both significantly upregulated, as well as the synthesis of glutamine involved in the reaction. Genes associated with nitrogen, phosphorus and sulfur transport were all upregulated to some extent **(Figure 5F)**. This is because these elements are essential for the replication and reproduction of phagosomes (35, 36). For example, the reduced availability of phosphorus in the growth environment limits the infection efficiency of cyanophage PP (37). The up-regulation of *pst* series genes can play a crucial role in assisting phage genome replication under phosphorus-depleted conditions (38).

### 3.6. Validation of the effects of cyanophage PP on 7942-PP via metabolomics

In order to verify the results and speculations of the transcriptomic analysis, metabolomics was carried out on 7942-sm and 7942-PP, and the contents of some key metabolites were detected (**Figure 6**). As shown in **Figure 4E** and **4F**, content of sucrose and glycogen decreased by 74% and 50% in 7942-PP compared to that in 7942-sm, respectively, demonstrating that the significant up-regulation of the *glk* gene resulted in their accelerated catabolism to Glucose 6-phosphate. From the metabolomic data, the levels of fructose 6-phosphate and fructose 1,6-bisphosphate were drastically reduced, whereas the level of glyceraldehyde 3-phosphate was virtually unchanged, which suggests that the cyanophage PP genome guides glucose 6-phosphate in PCC 7942 to participate more in the PPP and ED pathways and less in the EMP pathway. The large increase in the levels of glycerate 3-phosphate and phosphoenolpyruvate suggests that carbon metabolism in PCC 7942 is facilitated by the expression of the cyanophage PP genome. Although the promotion of PPP may lead to an increase in ribose 5-phosphate and ribulose 5-phosphate, the large amount of ribose 5-phosphate used for nucleotide and amino acid synthesis led to a decrease in the amount of ribose 5-phosphate and ribulose 5-phosphate in the experimental results. The ATP and NADPH contents of 7942-PP were greatly increased, which were 2.84 and 4.87 times higher than those of 7942-sm, respectively. This demonstrated that the genes of the cyanophage PP genome increased the intracellular ATP and NADPH content to meet the metabolic demand and energy supply of the additional phage genome replication by regulating the metabolism of cyanobacteria (39).

**Figure 6.**
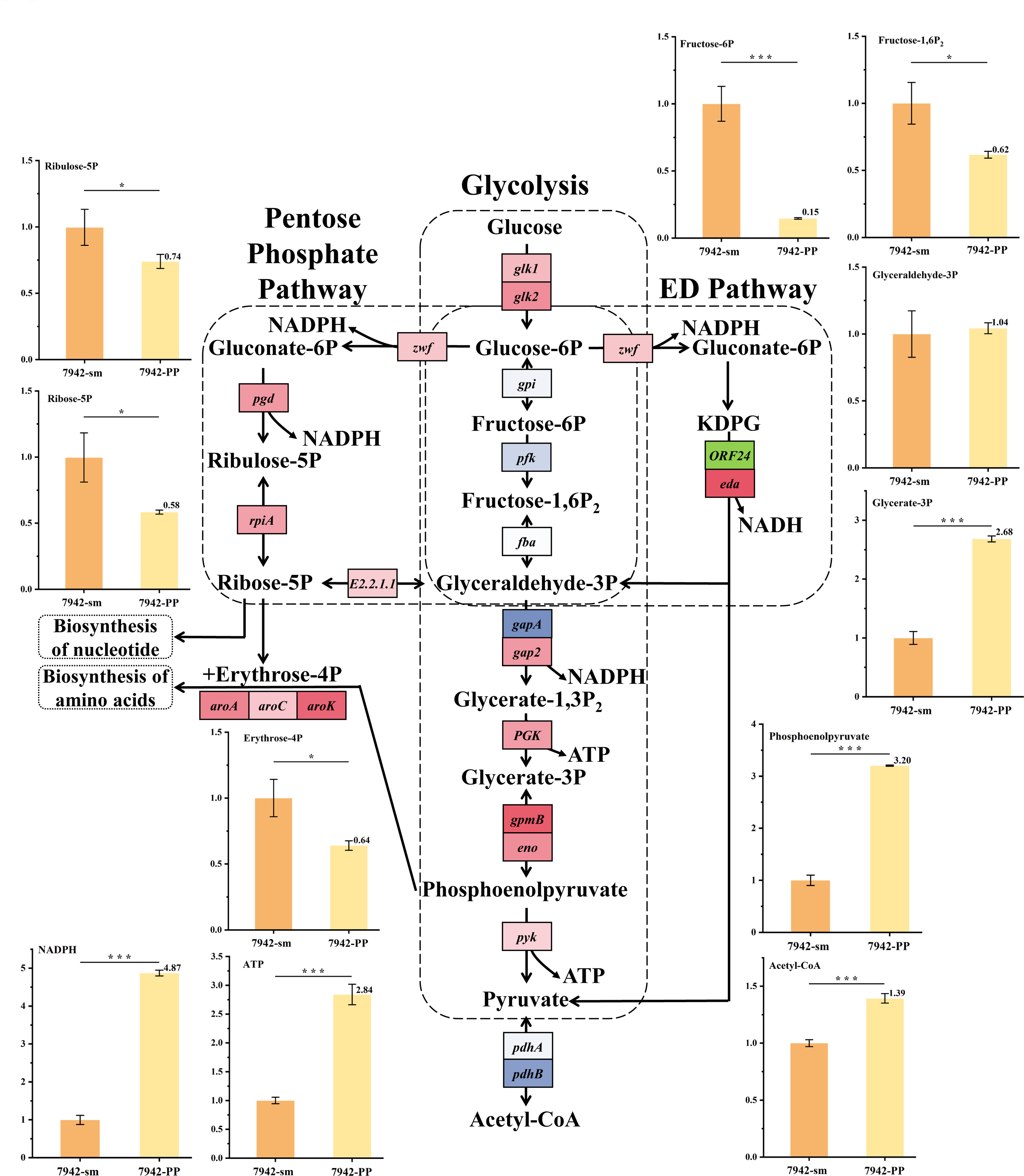
Changes in the content of metabolic components of the experimental strain 7942-PP. The value of 7942-sm is made to be 1 by the same proportion change, so that it is easy to observe the change of 7942-PP.

## 4. Discussion

Artificial design and assembly of cyanophage genomes with wide host range is a potential solution to solve the problem of cyanobacterial blooms, which is environmentally friendly and cost-effective. However, only a few freshwater cyanophage have been isolated and genomically sequenced so far. Previously, Hou et al. constructed a new synthetic *E. coli*-yeast shuttle vector that was able to transform an assembled portion of the phage PP genome (∼31kb) without toxicity genes into *E. coli* (40). Further, Chen et al. transformed the non-toxic phage genome PP4-8 portion into the model cyanobacterium *Synechocystis* sp. PCC 6803 (8), followed by the assembly of the full-length genome of cyanophage A-4L in yeast (16). However, the most critical step in rescuing cyanophages—the transformation of the full-length cyanophage genome, including toxicity genes, into cyanobacteria—has not yet been reported. This is primarily because transforming large genomes into cyanobacteria is generally only possible through conjugation transfer via *E. coli*. However, portions of the cyanophage genome are often toxic to *E. coli*, or non-toxic genes may co-express in a way that leads to *E. coli* death. In this study, we employ a tandem induction system using the LacI operator and theophylline riboswitch to control the genes toxic to E. coli, enabling their cloning in *E. coli*. We first integrated the longer portion of the genome into the cyanobacterial genome via triparental conjugative transfer through *E. coli*, obtaining an experimental strain. Subsequently, we integrated the remaining part of the genome, along with the toxicity genes controlled by dual induction, into this strain’s genome. Thus, we have successfully transformed the full-length genome of cyanophage PP, including terminal repeats and all toxicity genes, into cyanobacteria.

In general, the expression of the full-length genome of cyanophage PP retarded the growth of the heterologous host PCC 7942 and reduced its chlorophyll synthesis. From the results of transcriptome analysis, it was found that the transformation of the genome of cyanophage PP into PCC 7942 would have an impact on controlling the metabolism of the cyanobacterium to a certain extent in order to facilitate its own replication. In terms of the photosynthetic system, although the cyanophage PP genome reduces the synthesis of relevant photosynthetic proteins in cyanobacteria, it appears that efforts are made to maintain the photosynthetic capacity of the host by means such as increasing the efficiency of electron transfer, thus ensuring adequate energy supply (41). Energetically, CO_2_ fixation was inhibited while glucose catabolism was accelerated. The promotion of PPP, ED pathway and glycolysis of core module involving three-carbon compounds may all lead to the accumulation of ATP and NADPH within cyanobacteria, providing the conditions for cyanophage replication (39). In terms of proteins, the genome of cyanophage PP significantly reduces the synthesis of proteins that are abundant in cyanobacteria (e.g., Rubisco and a wide variety of photosynthetic proteins) while increasing the efficiency of the uptake of exogenous amino acids and the synthesis of specific amino acids *in vivo* in order to ensure that proteins required for phage replication are synthesized. In terms of nucleotides, the genome of cyanophage PP accelerated the transport of nitrogen, phosphorus, and sulfur by cyanobacteria, and accelerated the synthesis of PRPP and glutamine, which were further synthesized into nucleotides to satisfy the replication of the cyanophage genome.

It is possible that the cyanophage did not successfully replicate and lyse the cyanobacteria due to the inability of the heterologous host PCC 7942 to recognize the promoters of some genes in the cyanophage PP. To achieve the resurrection of artificial cyanophage genomes in future studies, the following methods can be considered: replacing the promoters in unexpressed genes with ones that PCC 7942 can recognize; transforming the full-length cyanophage PP genome into its hosts *P. boryanum* FACHB-240 using the methods described in this paper; and applying the approach of this paper to split the genomes of other cyanophage species into two parts and transform them into their corresponding host cyanobacteria.

## 5. Conclusion

In this study, we systematically screened and identified the potential toxic genes of the PP genome in *E. coli*, leading to the characterization of ORF6, ORF11, and ORF22. By controlling these three ORFs using tandem induction switches, PP1, PP2, and PP3 were successfully assembled and transferred into *E. coli*. Additionally, the full length of the PP genome was integrated into the genome of PCC 7942 via two rounds of homologous recombination. The gene expressions of the PP genome significantly influenced the photosynthesis and carbon fixation of PCC 7942, resembling the effects of a live cyanophage to some extent. To achieve the complete rescue of the artificial PP genome in the host, we believe that the development of genetic toolboxes in its original host, *P. boryanum* FACHB-240, is crucial for the future.

## Data Availability

The transcriptome data have been uploaded into NCBI with the accession number PRJNA1128005 (https://www.ncbi.nlm.nih.gov/sra/PRJNA1128005).

## Acknowledgements

This research was supported by grants from the National Key Research and Development Program of China (Grant no. 2018YFA0903000) and the National Natural Science Foundation of China (Grant no. 32371486).

## CRediT Authorship Contribution Statement

GL and JF carried out the main experiments and wrote the manuscript. XZ and YC helped analyze the data. TS and JJ conceived of the study and revised the manuscript. All authors read and approved the final manuscript.

## Supplementary Material

Supplementary material to this article can be found online.

