## Supplemental Tables and figures for "Expression and characterization of the complete cyanophage genome PP in the heterologous host *Synechococcus elongatus* PCC 7942"

**Supplementary Table S1.** List of the plasmids used or constructed in this study.

| **Plasmids** | **Description** |
| --- | --- |
| pRS426 | 2μ origin for *S. cerevisiae* and ColE1 (high-copy) for *E. coli*; Ura3 and Amp^R^ |
| pRSPP1 | pRS426 containing assembled fragements of PP1 from the PP genome |
| pRSPP2 | pRS426 containing assembled fragements of PP2 from the PP genome |
| pRSPP3 | pRS426 containing assembled fragements of PP3 from the PP genome |
| pRSPP4 | pRS426 containing assembled fragements of PP4 from the PP genome |
| pRSPP5 | pRS426 containing assembled fragements of PP5 from the PP genome |
| pRSPP6 | pRS426 containing assembled fragements of PP6 from the PP genome |
| pRSPP7 | pRS426 containing assembled fragements of PP7 from the PP genome |
| pRSPP8 | pRS426 containing assembled fragements of PP8 from the PP genome |
| pRSPP4-8 | pRS426 containing assembled fragements of PP4, PP5, PP6, PP7 and PP8 |
| pRSPP4-8+1 | PP1 was added to pRSPP4-8 |
| pRSPP4-8+2 | PP2 was added to pRSPP4-8 |
| pRSPP4-8+3 | PP3 was added to pRSPP4-8 |
| pRSPP1-F1 | pRS426 containing assembled partial fragements of PP1 |
| pRSPP1-F2 | pRS426 containing assembled partial fragements of PP1 |
| pRSPP1-F3 | pRS426 containing assembled partial fragements of PP1 |
| pRSPP1-F4 | pRS426 containing assembled partial fragements of PP1 |
| pRSPP2-F5 | pRS426 containing assembled partial fragements of PP2 |
| pRSPP2-F6 | pRS426 containing assembled partial fragements of PP2 |
| pRSPP2-F7 | pRS426 containing assembled partial fragements of PP2 |
| pRSPP3-F8 | pRS426 containing assembled partial fragements of PP3 |
| pRSPP3-F9 | pRS426 containing assembled partial fragements of PP3 |
| pRSPP1ΔORF5 | Deteling ΔORF5 of the PP1 fragments compared to pRSPP1 |
| pRSPP1ΔORF6 | Deteling ΔORF6 of the PP1 fragments compared to pRSPP1 |
| pRSPP1ΔORF7 | Deteling ΔORF7 of the PP1 fragments compared to pRSPP1 |
| pRSPP2ΔORF10 | Deteling ΔORF10 of the PP2 fragments compared to pRSPP2 |
| pRSPP2ΔORF11 | Deteling ΔORF11 of the PP2 fragments compared to pRSPP2 |
| pRSPP2ΔORF14 | Deteling ΔORF14 of the PP2 fragments compared to pRSPP2 |
| pRSPP3ΔORF22 | Deteling ΔORF22 of the PP3 fragments compared to pRSPP3 |
| pRSPP1-3Δ3 | pRS426 containing assembled fragements of PP1, PP2 and PP3 but without ORF6, ORF11 and ORF22 |
| pLS0 | CEN/ARS origin for *S. cerevisiae* and regulated origin (single copy) for *E. coli*; His3 and Cm^R^ |
| pLSPP1ΔORF5 | pLS0 containing assembled fragements of PP1 but without ORF5 from the PP genome |
| pLSPP1ΔORF6 | pLS0 containing assembled fragements of PP1 but without ORF6 from the PP genome |
| pLSPP1ΔORF7 | pLS0 containing assembled fragements of PP1 but without ORF7 from the PP genome |
| pLSPP2ΔORF10 | pLS0 containing assembled fragements of PP2 but without ORF10 from the PP genome |
| pLSPP2ΔORF11 | pLS0 containing assembled fragements of PP2 but without ORF11 from the PP genome |
| pLSPP2ΔORF14 | pLS0 containing assembled fragements of PP2 but without ORF14 from the PP genome |
| pLSPP3ΔORF22 | pLS0 containing assembled fragements of PP3 but without ORF22 from the PP genome |
| pLSPP1-3Δ3 | pLS0 containing assembled fragements of PP1, PP2 and PP3 but without ORF6, ORF11 and ORF22 |
| pLSPP1-3I3 | pLS0 containing assembled fragements of PP1, PP2 and PP3 with ORF6, ORF11 and ORF22 under the control of inducible switches |
| pLSPP4-8 | pLS0 containing assembled fragements of PP4, PP5, PP6, PP7 and PP8 |
| pLSPP1-8Δ3 | pLS0 containing assembled fragements of PP1, PP2, PP3, PP4, PP5, PP6, PP7 and PP8 but without ORF6, ORF11 and ORF22 |
| pLSPP1-8I3 | pLS0 containing assembled fragements of PP1, PP2 and PP3 with ORF6, ORF11 and ORF22 under the control of inducible switches |
| pSynPP4-8 | two homologous arms, λRed cassette and counter-selection marker *rpsL* was added to pLSPP4-8 |
| pSynPP1-3I3 | kanamycin-resistant cassette, Km^R^ and two homologous arms respectively targeting upstream of NSI site and partial of the PP4 was added to pLSPP1-3I3 |

**Supplementary Table S2**. Locations and functions of 41 predicted ORFs in the cyanophage PP genome

| GeneName | Start | End | KEGG | Function |
| --- | --- | --- | --- | --- |
| ORF1 | 419 | 685 | K02338 | DnaN, DNA polymerase sliding clamp subunit (PCNA homolog) |
| ORF2 | 774 | 1334 | K02338 | DnaN, DNA polymerase sliding clamp subunit (PCNA homolog) |
| ORF3 | 1331 | 1567 | - | hypothetical protein |
| ORF4 | 1545 | 1748 | - | hypothetical protein |
| ORF5 | 1949 | 3202 | K01952 | PurL, Phosphoribosylformylglycinamidine (FGAM) synthase |
| ORF6 | 3292 | 3837 | - | PurL, Phosphoribosylformylglycinamidine (FGAM) synthase |
| ORF7 | 3956 | 4624 | - | PurL, Phosphoribosylformylglycinamidine (FGAM) synthase |
| ORF8 | 4624 | 5133 | K00764 | PurF, Glutamine phosphoribosylpyrophosphate amidotransferase |
| ORF9 | 5078 | 5371 | K00764 | PurF, Glutamine phosphoribosylpyrophosphate amidotransferase |
| ORF10 | 5377 | 6636 | K00764 | PurF, Glutamine phosphoribosylpyrophosphate amidotransferase |
| ORF11 | 6641 | 8815 | - | Xanthine/uracil/vitamin C permease |
| ORF12 | 8876 | 9076 | K01940 | ArgG, Argininosuccinate synthase |
| ORF13 | 9089 | 9295 | K01940 | ArgG, Argininosuccinate synthase |
| ORF14 | 9299 | 11167 | K01940 | ArgG, Argininosuccinate synthase |
| ORF15 | 11149 | 11304 | - | hypothetical protein |
| ORF16 | 11497 | 11742 | K02990 | hypothetical protein |
| ORF17 | 11826 | 12788 | K02990 | RpsF, Ribosomal protein S6 |
| ORF18 | 13017 | 13193 | - | hypothetical protein |
| ORF19 | 13272 | 13514 | - | hypothetical protein |
| ORF20 | 13522 | 14025 | - | PiuC, Uncharacterized iron-regulated protein |
| ORF21 | 14038 | 14247 | - | PiuC, Uncharacterized iron-regulated protein |
| ORF22 | 14210 | 14548 | - | hypothetical protein |
| ORF23 | 14627 | 15487 | - | hypothetical protein |
| ORF24 | 15551 | 16753 | K01625 | Eda, 2-keto-3-deoxy-6-phosphogluconate aldolase |
| ORF25 | 16740 | 20201 | K00392 | GlyS, Glycyl-tRNA synthetase, beta subunit |
| ORF26 | 20299 | 25437 | - | ChaA, Ca2+/H+ antiporter |
| ORF27 | 25485 | 28469 | K00652 | Tas, Predicted oxidoreductases |
| ORF28 | 28471 | 29661 | K00652 | BioF, 7-keto-8-aminopelargonate synthetase and related enzymes |
| ORF29 | 29665 | 32748 | K00833 | BioA, Adenosylmethionine-8-amino-7-oxononanoate aminotransferase |
| ORF30 | 32738 | 33103 | - | Uncharacterized conserved protein |
| ORF31 | 33108 | 33773 | - | Predicted nucleotidyltransferases |
| ORF32 | 34050 | 34166 | - | hypothetical protein |
| ORF33 | 34216 | 35361 | - | major head protein; Phage capsid protein |
| ORF34 | 35384 | 36067 | - | head scaffolding protein |
| ORF35 | 36072 | 38027 | K00033 | Gnd, 6-phosphogluconate dehydrogenase |
| ORF36 | 38042 | 38467 | K00033 | Gnd, 6-phosphogluconate dehydrogenase |
| ORF37 | 38504 | 40201 | K01633 | FolB, Dihydroneopterin aldolase |
| ORF38 | 40307 | 40807 | - | ArsB, Na+/H+ antiporter NhaD and related arsenite permeases |
| ORF39 | 40813 | 41298 | K02902 | RpmB, Ribosomal protein L28 |
| ORF40 | 41319 | 41594 | - | hypothetical protein |
| ORF41 | 41599 | 41850 | - | hypothetical protein |

**Supplementary Table S3.** Expression of each predicted ORF in the cyanophage PP genome

| PP ORF | Gene Sizes | 1-1  Count | 1-1  FPKM | 1-2  Count | 1-2  FPKM | 1-3  Count | 1-3  FPKM | Start | End |
| --- | --- | --- | --- | --- | --- | --- | --- | --- | --- |
| ORF1 | 267 | 0 | 0 | 0 | 0 | 0 | 0 | 419 | 685 |
| ORF2 | 561 | 53 | 17.52 | 51 | 18.12 | 22 | 8.88 | 774 | 1334 |
| ORF3 | 237 | 0 | 0 | 0 | 0 | 0 | 0 | 1331 | 1567 |
| ORF4 | 204 | 0 | 0 | 0 | 0 | 0 | 0 | 1545 | 1748 |
| ORF5 | 1254 | 121 | 17.9 | 141 | 22.42 | 58 | 10.47 | 1949 | 3202 |
| ORF6 | 546 | 0 | 0 | 0 | 0 | 0 | 0 | 3292 | 3837 |
| ORF7 | 669 | 32 | 8.87 | 46 | 13.71 | 25 | 8.46 | 3956 | 4624 |
| ORF8 | 510 | 0 | 0 | 0 | 0 | 0 | 0 | 4624 | 5133 |
| ORF9 | 294 | 0 | 0 | 0 | 0 | 0 | 0 | 5078 | 5371 |
| ORF10 | 1260 | 720 | 105.98 | 718 | 113.61 | 294 | 52.83 | 5377 | 6636 |
| ORF11 | 2175 | 67 | 5.71 | 66 | 6.05 | 80 | 8.33 | 6641 | 8815 |
| ORF12 | 201 | 1 | 0.92 | 0 | 0 | 2 | 2.25 | 8876 | 9076 |
| ORF13 | 207 | 0 | 0 | 0 | 0 | 0 | 0 | 9089 | 9295 |
| ORF14 | 1869 | 138 | 13.69 | 99 | 10.56 | 44 | 5.33 | 9299 | 11167 |
| ORF15 | 156 | 6 | 7.13 | 3 | 3.83 | 11 | 15.96 | 11149 | 11304 |
| ORF16 | 246 | 28 | 21.11 | 11 | 8.91 | 5 | 4.6 | 11497 | 11742 |
| ORF17 | 963 | 305 | 58.74 | 200 | 41.41 | 89 | 20.92 | 11826 | 12788 |
| ORF18 | 177 | 14 | 14.67 | 15 | 16.9 | 8 | 10.23 | 13017 | 13193 |
| ORF19 | 243 | 52 | 39.69 | 38 | 31.18 | 28 | 26.09 | 13272 | 13514 |
| ORF20 | 504 | 25 | 9.2 | 23 | 9.1 | 12 | 5.39 | 13522 | 14025 |
| ORF21 | 210 | 1 | 0.88 | 0 | 0 | 0 | 0 | 14038 | 14247 |
| ORF22 | 339 | 8 | 4.38 | 10 | 5.88 | 6 | 4.01 | 14210 | 14548 |
| ORF23 | 861 | 29 | 6.25 | 17 | 3.94 | 71 | 18.67 | 14627 | 15487 |
| ORF24 | 1203 | 94 | 14.49 | 117 | 19.39 | 45 | 8.47 | 15551 | 16753 |
| ORF25 | 3462 | 44 | 2.36 | 42 | 2.42 | 83 | 5.43 | 16740 | 20201 |
| ORF26 | 5139 | 150 | 5.41 | 179 | 6.94 | 46 | 2.03 | 20299 | 25437 |
| ORF27 | 2985 | 90 | 5.59 | 104 | 6.95 | 47 | 3.56 | 25485 | 28469 |
| ORF28 | 1191 | 17 | 2.65 | 28 | 4.69 | 8 | 1.52 | 28471 | 29661 |
| ORF29 | 3084 | 32 | 1.92 | 35 | 2.26 | 24 | 1.76 | 29665 | 32748 |
| ORF30 | 366 | 5 | 2.53 | 3 | 1.63 | 3 | 1.86 | 32738 | 33103 |
| ORF31 | 666 | 48 | 13.37 | 32 | 9.58 | 50 | 17 | 33108 | 33773 |
| ORF32 | 117 | 0 | 0 | 1 | 1.7 | 3 | 5.8 | 34050 | 34166 |
| ORF33 | 1146 | 41 | 6.64 | 34 | 5.91 | 125 | 24.69 | 34216 | 35361 |
| ORF34 | 684 | 14 | 3.8 | 7 | 2.04 | 8 | 2.65 | 35384 | 36067 |
| ORF35 | 1956 | 21 | 1.99 | 22 | 2.24 | 10 | 1.16 | 36072 | 38027 |
| ORF36 | 426 | 1 | 0.44 | 3 | 1.4 | 3 | 1.59 | 38042 | 38467 |
| ORF37 | 1698 | 49 | 5.35 | 38 | 4.46 | 32 | 4.27 | 38504 | 40201 |
| ORF38 | 501 | 27 | 9.99 | 24 | 9.55 | 7 | 3.16 | 40307 | 40807 |
| ORF39 | 486 | 1888 | 720.47 | 1679 | 688.76 | 115 | 53.57 | 40813 | 41298 |
| ORF40 | 276 | 8 | 5.38 | 7 | 5.06 | 4 | 3.28 | 41319 | 41594 |
| ORF41 | 252 | 20 | 14.72 | 22 | 17.41 | 16 | 14.37 | 41599 | 41850 |

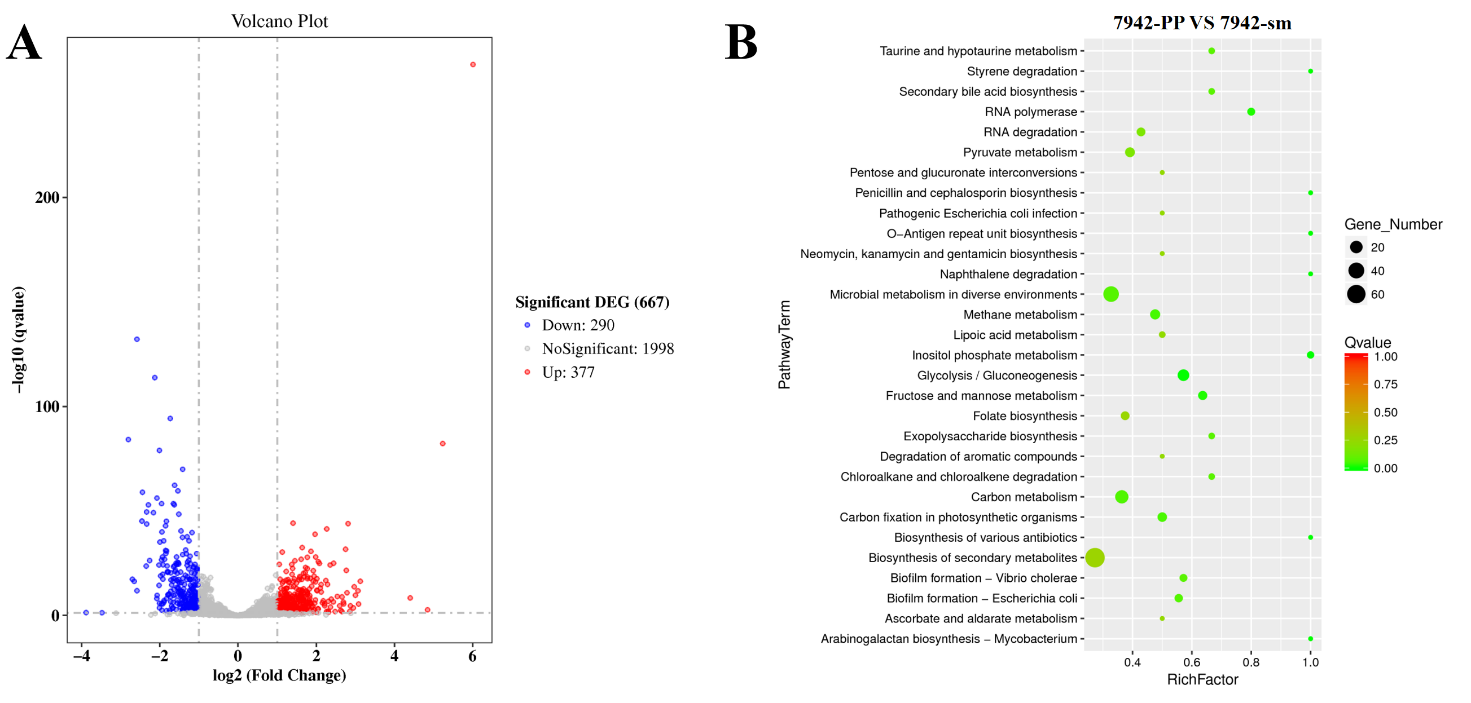

**Figure S1**. Up and down regulated genes (A) and their functional categories obtained from the transcriptomic data.
